# Specialized DNA structures act as genomic beacons for integration by evolutionarily diverse retroviruses

**DOI:** 10.1101/2020.02.23.959932

**Authors:** Hinissan P. Kohio, Hannah O. Ajoge, Macon D. Coleman, Emmanuel Ndashimye, Richard M. Gibson, Eric J. Arts, Stephen D. Barr

## Abstract

Retroviral integration site targeting is not random and plays a critical role in expression and long-term survival of the integrated provirus. To better understand the genomic environment surrounding retroviral integration sites, we performed an extensive comparative analysis of new and previously published integration site data from evolutionarily diverse retroviruses from seven genera, including different HIV-1 subtypes. We showed that evolutionarily divergent retroviruses exhibited distinct integration site profiles with strong preferences for non-canonical B-form DNA (non-B DNA). Whereas all lentiviruses and most retroviruses integrate within or near genes and non-B DNA, MMTV and ERV integration sites were highly enriched in heterochromatin and transcription-silencing non-B DNA features (e.g. G4, triplex and Z-DNA). Compared to *in vitro*-derived HIV-1 integration sites, *in vivo*-derived sites are significantly more enriched in transcriptionally silent regions of the genome and transcription-silencing non-B DNA features. Integration sites from individuals infected with HIV-1 subtype A, C or D viruses exhibited different preferences for non-B DNA and were more enriched in transcriptionally active regions of the genome compared to subtype B virus. In addition, we identified several integration site hotspots shared between different HIV-1 subtypes with specific non-B DNA sequence motifs present at these hotspots. Together, these data highlight important similarities and differences in retroviral integration site targeting and provides new insight into how retroviruses integrate into genomes for long-term survival.

**Graphical Abstract:** 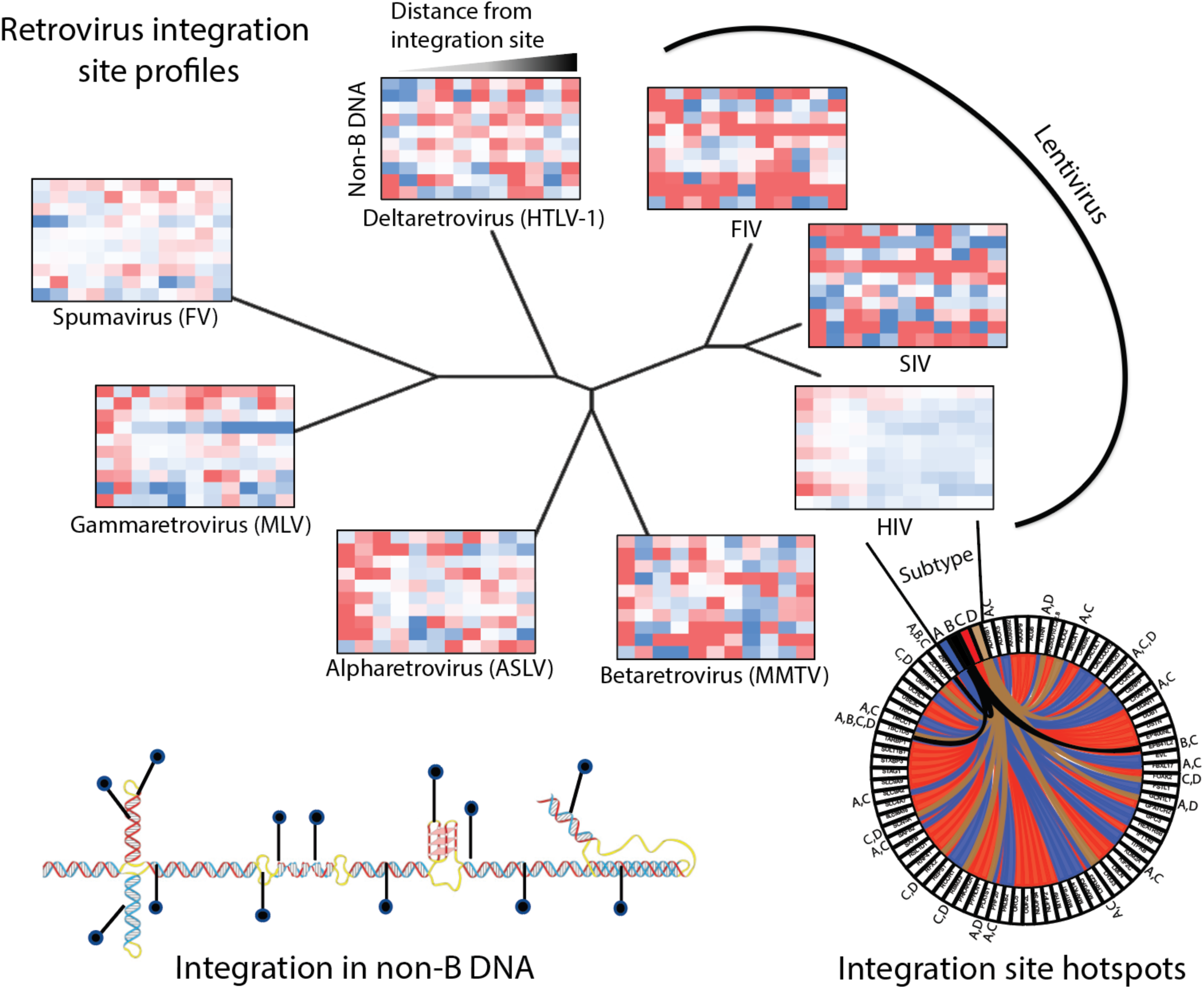

Schematic comparing integration site profiles from evolutionarily diverse retroviruses. Upper left, heatmaps showing the fold-enrichment (blue) and fold-depletion (red) of integration sites near non-B DNA features (lower left). Lower right, circa plot showing integration site hotspots shared between HIV-1 subtype A, B, C and D virus.

## INTRODUCTION

Retroviruses are divided into seven genera *alpha-, beta-, gamma-, delta-*, and *epsilon-Retroviridae, Spumaviridae*, and *Lentiviridae*). Following entry of a retrovirus into cells, the viral RNA genome is converted into double-stranded complimentary DNA (cDNA). In the cytoplasm, the cDNA associates with several viral and host proteins to form a pre-integration complex (PIC). Soon after completion of cDNA synthesis, the viral DNA ends are primed by the enzyme integrase in a process called 3’ processing. After docking with a host chromosome, the viral cDNA undergoes a strand transfer reaction resulting in the insertion of the retroviral genome into the chromosome. Host repair enzymes are then thought to complete integration (*1*). Insertion of the retroviral genome results in a persistent life-long infection. Selection of integration sites in the genome by the PIC is not random. The gamma- and deltaretroviruses (e.g. murine leukemia virus (MLV), human T cell leukemia virus Type 1 (HTLV-1)) and foamy virus (FV) favor integration around *transcription start sites* (*TSS*) (*2*–*4*). Alpharetroviruses (e.g. avian sarcoma leukosis virus (ASLV)) and human endogenous retroviruses (ERVs) show no strong preferences, with integration only slightly favored in transcription units or the 5’ end of genes (*5*–*8*).

Lentiviruses such as human immunodeficiency virus type 1 (HIV-1), simian immunodeficiency virus (isolated from a pig-tailed macaque) (SIV_mne_) and feline immunodeficiency virus (FIV) strongly favor integration in active transcription units (*9*–*11*). HIV integrations sites are also associated with regions of high G/C content, high gene density, short introns, high frequencies of short interspersed nuclear elements (*SINEs*) (e.g. Alu repeats), low frequencies of long interspersed nuclear elements (*LINEs*), and characteristic epigenetic modifications (*9, 12*). Thus far, integration site analyses have only been conducted on HIV-1 subtype B infections. Based on phylogenetic analyses of full-length genomic sequences, HIV-1 isolates are classified into four distinct groups: group M, N, O and P (*13*). HIV-1 M group accounts for the majority of the global pandemic and is subdivided into nine subtypes or clades (A, B, C, D, F, G, H, J and K). Additionally, several circulating recombinant forms (CRF) and unique recombinant forms (URF) have been identified, which are the result of a recombination event between two or more different subtypes. HIV-1 geographical prevalence is extremely diverse. Subtype C represents more than 50% of the infection worldwide and is prevalent in Africa and Asia. Subtype B infection is common in the Americas, Europe, Australia, and part of South Asia, Northern Africa and the Middle East. Subtypes A, D, F, G, H, J and K occur mostly in Sub-Saharan Africa, whereas infections with groups N, O and P have been found in confined regions of West-Central Africa.

Several models, not mutually exclusive, have been proposed to explain integration site selection. In the chromatin accessibility model, the structure of chromatin influences accessibility of target DNA sequences to PICs. *In vivo* target DNA is not expected to be naked but rather wrapped in nucleosomes. Wrapping target DNA in nucleosomes does not reduce integration, but instead creates hotspots for integration at sites of probable DNA distortion (*14, 15*). Distortion of DNA in several other protein-DNA complexes has also been shown to favour integration in the major grooves facing outwards from the nucleosome core (*16, 17*). Although chromatin structure can facilitate integration, chromatin accessibility cannot solely explain the differences observed in integration site preferences.

The protein tethering model suggests that a cellular protein, potentially specific for each retroviral genera, act as tethering factors between chromatin and the PIC. The most characterized tethering factor identified to date is lens epithelium-derived growth factor and co-factor p75 (LEDGF/p75) (also known as PSIP1/p75) (*18*–*20*). LEDGF/p75 interacts with HIV integrase and tethers the PIC to genomic DNA in transcriptionally active genes marked by specific histone modifications such as H3K20me1, H3K27me1 and H3K36me3. In similar fashion, host protein bromodomain/extraterminal domain proteins (BETs) bound to acetylated histones (e.g. H327ac and H3K9ac) interact with MLV integrase and tethers the PIC to genomic DNA in transcriptionally active promoters, enhancers and super enhancers (*21*–*24*). LEDGF/p75 and BET depletion studies demonstrated that integration still occurs but with reduced efficiency and an altered integration site selection profile (*20, 22*–*28*). Several cellular proteins have been proposed to facilitate integration including barrier to autointegration factor (BAF), high mobility group A1 (HMGA1), integrase interactor 1 (Ini-1), and heat shock protein 60 (Hsp60) (*29*–*31*). Some of these proteins might contribute to PIC function by coating and condensing the viral DNA, thereby assisting the assembly of the viral nucleoprotein complexes. Several cellular chromatin proteins have also been suggested to influence integration such as Ini-1, EED, SUV39H1, and HP1γ (reviewed in references (*12, 32, 33*)). Cleavage and polyadenylation specificity factor 6 (CPSF6) facilitates HIV-1 PIC import and helps direct the PIC to the nuclear interior to locate gene-dense euchromatin for integration (*34*–*38*). Recently, Apolipoprotein B Editing Complex (APOBEC3) was identified as a potential new host factor that also influences integration site selection by promoting a more transcriptionally silent integration site profile (*39*). Moreover, just as DNA-binding proteins can promote integration, they can also block access of integration complexes, creating regions refractory for integration (*16, 40, 41*). Binding to different factors or differential binding affinity to a factor by the PIC could modulate integration site selection between lentivirus types or even between different HIV-1 subtypes.

Previous analyses of primary DNA sequences (∼20 base pairs (bp)) flanking integration sites only revealed a weak consensus motif (*42*). We previously assessed a broader window of primary sequence (80 bp) around HIV-1 integration sites discovered that HIV-1 integration sites were highly enriched near specialized genomic features called non-B DNA (*43*). Non-B DNA form functional secondary structures in our genome formed by specific nucleotide sequences that exhibit non-canonical DNA base pairing. At least 10 non-B DNA conformations are identified including inverted repeats, direct repeats, mirror repeats, short-tandem repeats, guanine-quadruplex (G4), A-phased, cruciform, slipped, triplex and Z-DNA (*44, 45*). Here we present a comparative analysis of integration site profiles of evolutionarily diverse retroviruses and identify striking similarities and differences among all retroviruses, especially towards non-B DNA.

## MATERIAL AND METHODS

### Ugandan and Zimbabwean cohort description

Details pertaining to the Uganda study population have been reported previously (*46*–*49*). Briefly, women who became HIV infected while participating in the Hormonal Contraception and Risk of HIV Acquisition Study in Uganda were enrolled upon primary infection with HIV-1 into a subsequent study, the Hormonal Contraception and HIV-1 Genital Shedding and Disease Progression among Women with Primary HIV Infection (GS) Study. Ethical approval was obtained from the Institutional Review boards (IRBs) from the Joint Clinical Research Centre and UNST in Uganda, from University of Zimbabwe, from the University Hospitals of Cleveland, and recently, from Western University. All adult subjects provided written informed consent and no child participants were included in the study. Protocol numbers and documentation of these approvals/renewals are available upon request. Blood and cervical samples were collected every month for the first six months, then every three months for the first two years, and then every six months up to 9.5 years. Women who had CD4 lymphocyte counts of 200 cells/ml and/or who developed severe symptoms of HIV infection (WHO clinical stage IV or advanced stage III disease) were offered combination antiretroviral therapy (cART) and trimethoprim-sulfamethoxazole (for prophylaxis against bacterial infections and *Pneumocystis jeroveci* pneumonia).

### HIV-1 integration site library

Total genomic DNA was extracted from PBMCs using the QIAmp DNA mini kit (Qiagen) and processed for integration site analysis and sequenced using the Illumina MiSeq platform as previously described (*43, 50*). Samples were sequenced at the London Regional Genomics Centre/Robarts Research Institute (Western University, Canada) and Case Western Reserve University (USA).

### Computational analysis

Fastq sequencing reads were quality trimmed and unique integration sites identified using our in-house bioinformatics pipeline Barr Lab Integration Site Identification Pipeline (BLISIP version 2.9) and included the following updates: bedtools (v2.25.0), bioawk (awk version 20110810), bowtie2 (version 2.3.4.1), and restrSiteUtils (v1.2.9) (*39, 43*). All non-B DNA motifs were defined according to previously established criteria (*51*). The G-quadruplex algorithm identifies four or more individual G-runs of at least three nucleotides in length. The algorithm requires at least one nucleotide between each run and considers up to seven nucleotides as spacer, including guanines (*51*). Lamina-associated domains (*LADs*) were retrieved from http://dx.doi.org/10.1038/nature06947 (*52*). HIV-1 LTR-containing fastq sequences were identified and filtered by allowing up to a maximum of five mismatches with the reference NL4-3 LTR sequence and accepted if the LTR sequence had no match with any region of the human genome (GRCh37/hg19). Matched random control integration sites (28,800 in total) were generated by matching each experimentally determined site with 50 random sites *in silico* that were constructed to be the same number of bases from the restriction site as was the experimental site, as previously described (*53*). All genomic sites in each dataset that hosted two or more sites (i.e. identical sites) were collapsed into one unique site for the analysis. Sites that could not be unambiguously mapped to a single region in the genome were excluded from the study. Integration site profile heatmaps were generated using our in-house python program BLISIP heatmap (BHmap v1.0).

### Data and software availability

The sequences reported in this paper will be deposited in the National Center for Biotechnology Information Sequence Read Archive (NCBI SRA) upon acceptance for publication.

## RESULTS

### Integration site dataset acquisition and analyses

Previous integration site analyses of evolutionarily diverse retroviruses identified three general integration site profiles in which retroviruses can be grouped. For example, HIV-1, SIV_mne_ and FIV integrate into transcriptionally active genes; MLV, HTLV-1 and FV integrate near *TSS*; ASLV, HTLV and MMTV show little preference for any genomic feature. The integration site profiles of these evolutionarily diverse retroviruses with respect to non-B DNA was previously unknown. To determine integration site profiles of these viruses, including *endogenous retroviruses* (*ERVs*) present within the human genome, we analyzed ∼169,910 integration sites from previously published datasets (**Table S1**). To allow for a more comparable analysis of integration site profiles, we updated these integration site profiles using the human genome assembly GRCh37/19. Our update also included an assessment of genomic features not included in some of the previous retroviral integration site studies. We used our previously developed in-house bioinformatics pipeline called the Barr Lab Integration Site Pipeline (BLISIP) to generate integration site profiles from the different retroviral integration site datasets (*39, 43, 50*). BLISIP measures integration site enrichment near the genomic features CpG islands, *DNAseI hypersensitivity sites* (*DHS*), *ERVs*, heterochromatic DNA regions (e.g. *lamin-associated domains* (*LADs*) and satellite DNA), *SINEs* and *LINEs* respectively, *low complexity repeats* (*LCRs*), oncogenes, genes, simple repeats and *TSS*. In addition, BLISIP measures enrichment near the non-B DNA features A-phased motifs, cruciform motifs, direct repeats, G4 motifs, inverted repeats, mirror repeats, short tandem repeats, slipped motifs, triplex motifs and Z-DNA motifs. Our analyses focused on unique integration sites and excluded sites arising from clonal expansion, sites falling in repeat regions that could not be uniquely identified, and regions that could not be confidently placed on a specific chromosome (e.g. ChrUn). Enrichment of integration sites within genomic features was determined by comparing the proportion of sites with either a matched random control (MRC) to account for restriction site bias in the cloning procedure during library construction, or a random control (RC) for comparison of datasets that used DNA shearing/fragmentation during library construction (**Table S1**).

### Evolutionarily divergent retroviruses exhibit distinct integration site profiles

Integration sites were quantified and placed in five bins based on their distance from each genomic feature (within the feature, 1-499 bp, 500-4,999 bp, 5,000-49,999 bp, and >49,000 bp). Heatmaps from each retrovirus showing the fold-enrichment and fold-depletion in each bin were compared to MRC or RC (**Figure 1A**). Consistent with previous studies (*9*–*11*), HIV-1, SIV_mne_ and FIV integration sites are significantly enriched within genes (64%, 84%, 90% respectively) (*P* < 0.0001) **(Figure 1A and 1B, Table S2)**. Our analysis of HTLV-1, ASLV and MLV also agreed with previous reports, confirming that these viruses exhibit only modest preferences for integration within *genes* with 54%, 56% and 56% of integration sites found within *genes* respectively **(Figure 1A and 1B, Table S2)** (*54, 55*). In contrast, MMTV, FV and ERVs showed no preference for integration into genes (47%, 43% and 32% respectively). No retrovirus showed a preference for integration directly into *TSS*; however, MLV, HTLV-1 and FV showed significant enrichment of integration sites near *TSS* and CpG islands (*P* <0.0001) **(Figure 1A, Table S2)**. Repetitive elements, such as *LINEs, SINEs, ERVs* (e.g. retrotransposons), satellite DNA, simple repeats (e.g. microsatellites), and *LCRs* account for nearly half of the human genome sequence. However, no strong preference for integration into these regions was observed for any of the retroviruses except for HIV-1 and FV which targeted *SINEs*, and FV, MMTV and HTLV-1 which targeted satellite DNA **(Figure 1A).** Pairwise analyses of the different integration site profile preferences (fold enrichment and depletion values) showed that HIV-1, MMTV and FIV had the most similar integration site preferences for genomic features overall, whereas SIV, MLV and ERVs had the least similarity to the other retroviruses (**Figure 1C**).

**Figure 1:**
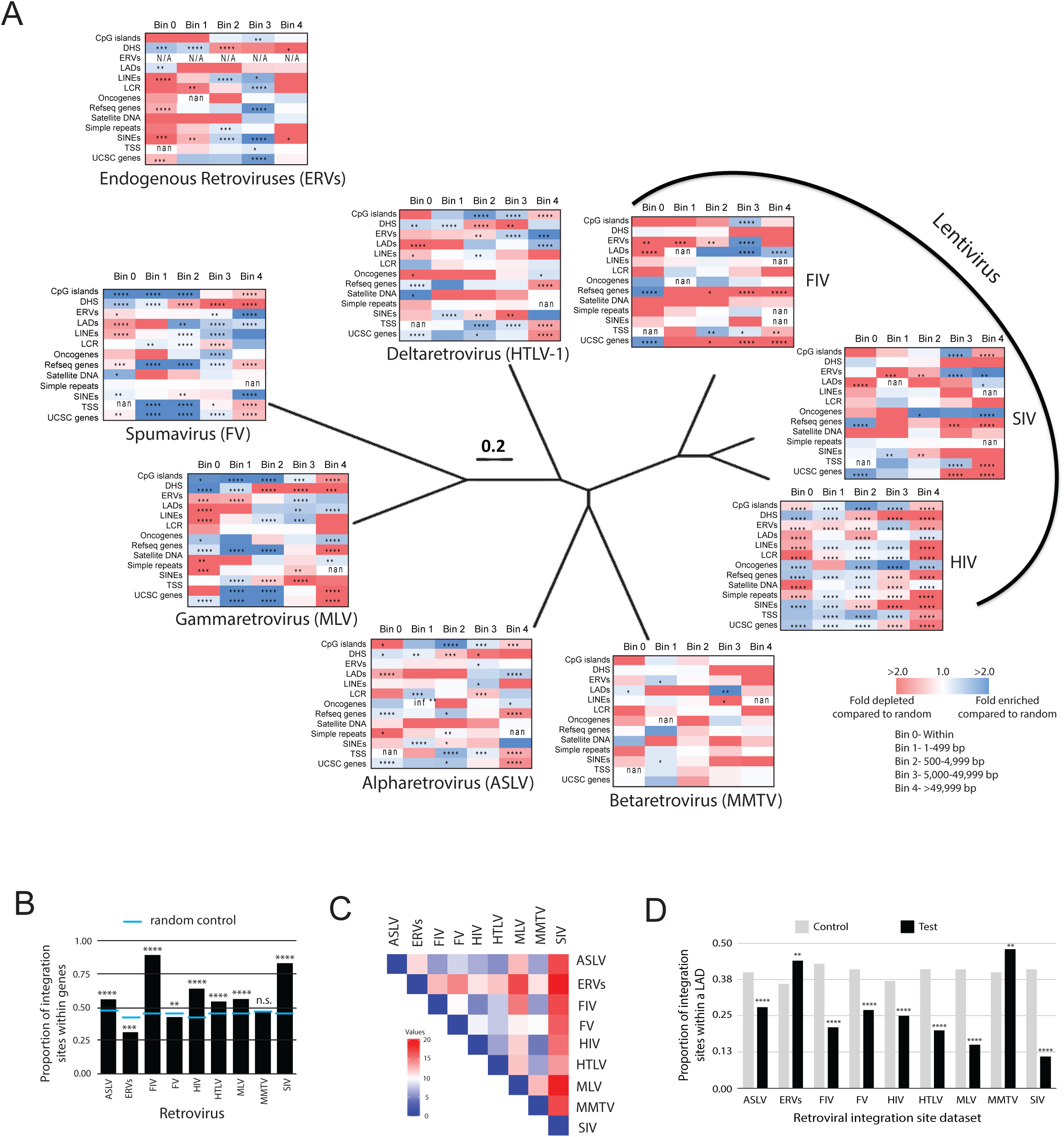
Evolutionarily diverse retroviruses exhibit distinct integration site preferences. **A**, Heatmaps depicting the fold enrichment or depletion of integration sites near common genomic features compared to matched random controls. Darker shades represent higher fold-changes in the ratio of integration sites to matched random control sites. Bins represent the distance of the integration sites from each genomic feature. Bin 0 = within the feature; Bin 1= 1-499 bp; Bin 2 = 500-4,999 bp; Bin 3 = 5,000-49,999 bp; Bin 4 = >49,999 bp. Heatmaps of the diverse retrovirus genera were superimposed on a BioNJ tree constructed using their reverse transcriptase amino acid sequences using the Dayhoff substitution model with 1000 bootstraps. All branches are scaled according to number of amino acids changes per site. The phylogenetic tree shows the evolutionary relatedness of the different retrovirus genera only. Significant differences are denoted by asterisks (*P < 0.05; **P < 0.01; ***P < 0.001; ****P < 0.0001) (Fisher’s Exact test). **B**, Proportion of retroviral integration sites located within genes compared to the random control (blue line). **C**, Pairwise analysis was performed on the retroviral integration site profile preferences (fold enrichment and depletion values) using Euclidean distance as the measurement method (Heatmapper, Babicki et al., 2016). Stronger relationships between retroviral integration site profiles are indicated by darker blue color in the pairwise distance matrix. **D**, Nuclear localization of integration sites were determined by quantifying the proportion of total integrations that fell within a lamin-associated domain (LAD) (=1) as opposed to outside a LAD (=0). *P < 0.05; **P < 0.01; ***P < 0.001; ****P < 0.0001; Fisher’s Exact test. Infinite number (inf), 1 or more integrations were observed when 0 integrations were expected by chance. Not a number (nan), 0 integrations were observed and 0 were expected by chance. HIV-1 = human immunodeficiency virus, SIV_mne_ = simian immunodeficiency virus (isolated from a pig-tailed macaque), FIV= feline immunodeficiency virus, HTLV-1= human T-lymphotrophic virus Type 1, MLV = murine leukemia virus, FV = foamy virus, ASLV = avian sarcoma leucosis virus, MMTV = mouse mammary tumor virus and ERVs = endogenous retroviruses.

### MMTV and ERVs strongly favour integration into heterochromatin

The nuclear architecture influences HIV-1 integration site selection and proviral expression (*52, 56*). HIV-1 strongly disfavors integration into heterochromatin positioned in *LADs* at the nuclear periphery, although some integration does occur in these regions, contributing to the latent reservoir (*57*). We asked whether other retroviruses similarly disfavor integration into *LADs*. Like HIV-1, most retroviruses significantly disfavored integration into *LADs* with only 11-28% of sites falling within *LADs* **(Figure 1D, Table S2).** In contrast, 48% of MMTV and 44% of ERV integration sites were significantly enriched in *LADs*.

### Evolutionarily divergent retroviruses target non-B DNA for integration

Non-B DNA is a genomic correlate of HIV-1 integration site selection, but its influence on integration site targeting of other retroviruses was previously unknown (*43*). Analysis of the different retroviral integration site profiles showed that all retroviruses exhibited enriched integration near non-B DNA (**Figure 2A and Table S3**). FV, HTLV-1 and SIV_mne_ were the only retroviruses with an enrichment of sites within 50 bp of multiple non-B DNA features, whereas other retroviruses tended to integrate more distal (100-500 bp) to the features. HIV-1 integration sites were enriched near all non-B DNA features except A-phased DNA. SIV_mne_ and FIV sites were enriched near inverted repeats, mirror repeats and short tandem repeats. HTLV-1 sites were enriched near A-phased, cruciform and inverted repeats. FV sites were enriched near G4 and Z-DNA. MLV sites were enriched near G4, triplex and Z-DNA. ASLV sites were enriched in slipped and triplex DNA. MMTV sites were enriched in short-tandem repeats, G4, slipped, triplex and Z-DNA. ERV sites were enriched near cruciform and triplex DNA. Pairwise analyses of the different integration site profiles showed that HIV-1 and FV had the most similar preferences for targeting non-B DNA, whereas SIV_mne_, ERVs and FIV had the least similarity to the other retroviruses (**Figure 2B**). Taken together, these data show that evolutionarily diverse retroviruses target non-B DNA for integration and that each virus exhibits distinct preferences for certain non-B DNA features.

**Figure 2:**
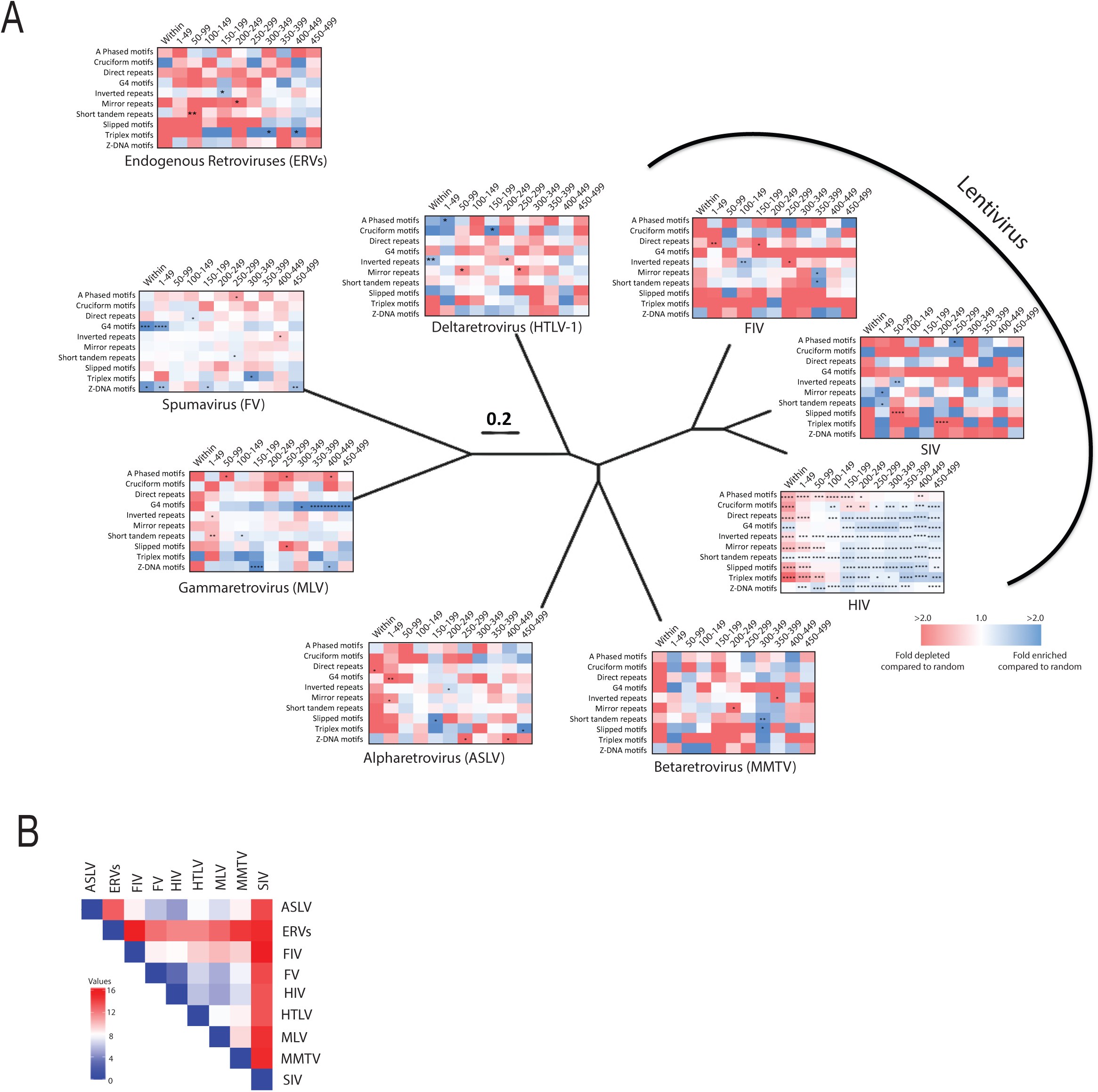
Evolutionarily diverse retroviruses target non-B DNA for integration. **A**, Heatmaps illustrating the fold-enrichment or depletion of unique retroviral integration sites near non-B DNA features compared to matched random controls. Darker shades represent higher fold-changes in the ratio of integration sites to matched random control sites. Significant differences are denoted by asterisks *P < 0.05; **P < 0.01; ***P < 0.001; ****P < 0.0001) (Fisher’s Exact test). **B**, Pairwise analysis was performed on the retroviral integration site profile preferences using Euclidean distance as the measurement method (Heatmapper, Babicki et al., 2016). Stronger relationships between retroviral integration site profiles are indicated by darker blue color in the pairwise distance matrix.

### HIV-1 Integration site profiles differ between *in vitro*- and *in vivo*-derived datasets

Despite a large number of integration sites from HIV-1 infected individuals, many of the previous integration site studies have been performed on infections done *in vitro* (*12, 58*). To determine if the HIV-1 integration site profiles from *in vitro*-derived infections differed from those from *in vivo*-derived infections, we analyzed and compared nine previously published HIV-1 integration site datasets from publicly available databases, totalling 13,601 *in vitro*-derived sites and 139,480 *in vivo*-derived sites **(Table S1)**. As expected, integration sites were significantly enriched in genes compared to the random controls in both the *in vitro-* and *in vivo*-derived datasets; however, only 62% of total *in vivo*-derived integration sites were in genes compared to 83% in the *in vitro* dataset (*P* < 0.0001) (**Figure 3A, 3B and Tables S4**). In contrast with the *in vitro* dataset, integration sites in the *in vivo*-derived dataset were highly enriched near most non-B DNA except A-phased DNA **(Figure 3C and Table S4)**. Notably, *in vivo*-derived sites were significantly more enriched near G4, triplex and Z-DNA than *in vitro*-derived sites **(Figure 3C, 3D and Table S4)**. Together, these data show that there are substantial differences in HIV-1 integration site targeting preferences between *in vitro*-derived and *in vivo*-derived datasets.

**Figure 3:**
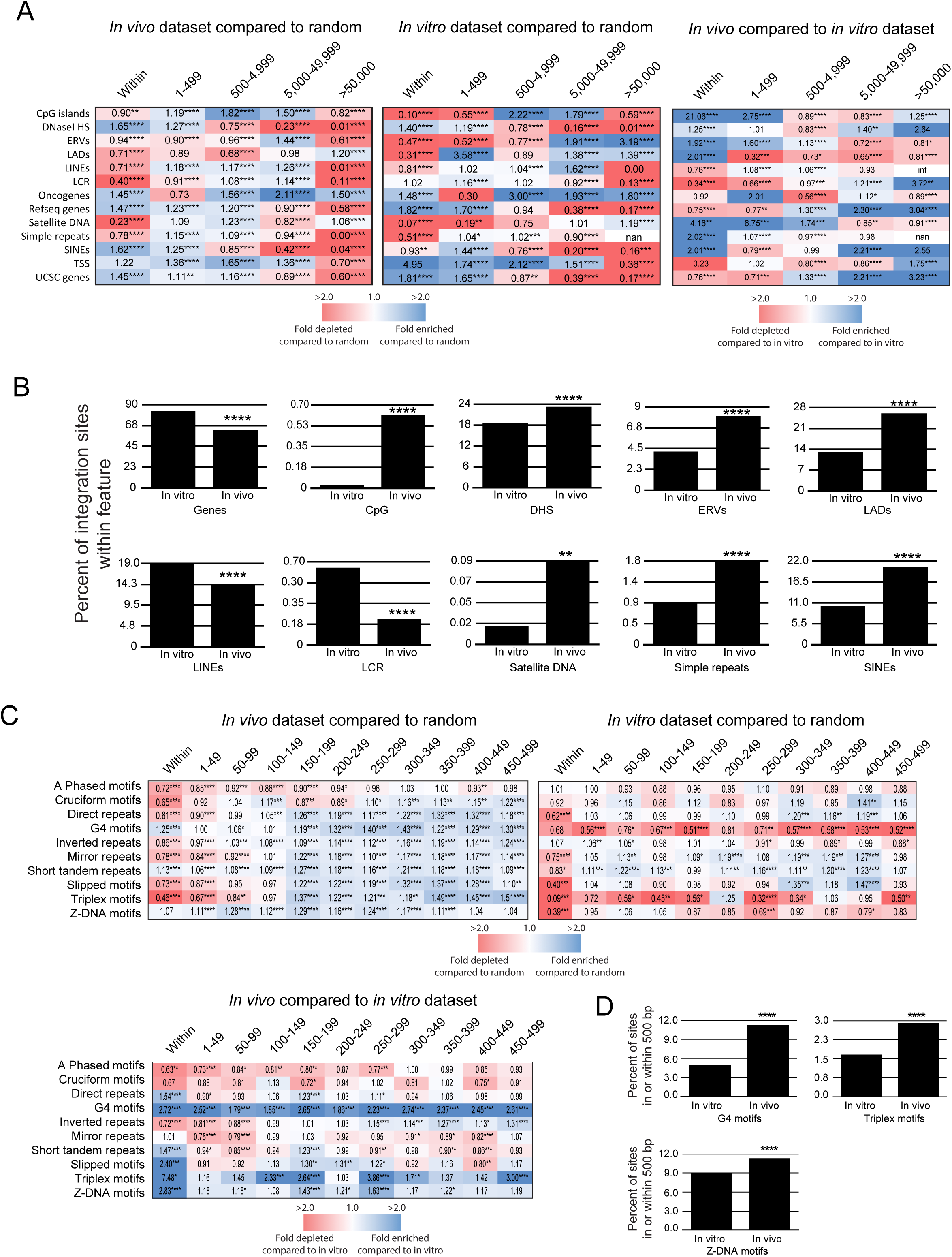
Integration site profiles differ between *in vitro*- and *in vivo*-derived datasets. **A**, Heatmaps illustrating the fold-enrichment or depletion of unique integration sites (compared to the matched random control) near common genomic features from *in vivo-*derived datasets (n= 167,098 sites) and *in* vitro-derived datasets (n= 13,601 sites). Inset numbers represent the fold-change in the percentage of integration sites. **B**, Comparison of the percentage of integration sites within 5,000 bp of common genomic features between *in vitro-* and *in vivo*-derived datasets. **C**, Heatmaps illustrating the fold-enrichment of unique integration sites compared to the matched random control near non-B DNA from *in vivo-* and *in vitro*-derived datasets. **D**, Comparisons of the percentage of integration sites within 500 bp of G4, triplex and Z-DNA between *in vitro-* and *in vivo*-derived datasets. Significant differences are denoted by asterisks *P < 0.05; **P < 0.01; ***P < 0.001; ****P < 0.0001) (Fisher’s Exact test).

### Integration site profiles differ in individuals infected with HIV-1 subtype A, B, C or D

Thus far, integration site profiles have been extensively analyzed for HIV-1 subtype B infections, which represents only ∼10% of the infections worldwide. We asked if the integration site profiles from individuals infected with HIV-1 non-subtype B virus were similar to those infected with subtype B virus. In the integrase enzyme, the amino acid difference between subtypes is as high as 16% (subtype B vs. C) but typically less than 8% in IN of HIV-1 isolates of a specific subtype. This level of amino acid diversity is the highest among the enzymes encoded by the HIV-1 *pol* gene. The greatest diversity within HIV-1 IN is found in the C-terminal domain (CTD), which is involved in genomic DNA binding. Thus, it is reasonable to propose that we would observe differences in selectivity for integrations in non-B form DNA. Genomic DNA was isolated from peripheral blood mononuclear cells (PBMCs) from a cohort of women in Uganda and Zimbabwe infected with HIV-1 subtype A, C or D and used to generate integration site libraries. Integration site profiles were generated from a total of 48 infected individuals (16 subtype A, 19 subtype C and 13 subtype D) and compared to the integration site profile from 14 individuals infected with subtype B virus generated from previously published datasets **(Table S5)** (*59, 60*).

As previously observed with HIV-1 subtype B infections, integration sites from all HIV-1 subtype viruses were enriched in genes (**Figure 4A and 4B and Table S6**). Notably, subtypes A, C and D had significantly more integration sites in genes compared to subtype B (A:82%, B:63%, C:71%, D:78%) (**Figure 4C**). Integration sites for all subtypes were enriched near *DHS*; however, subtypes A, C and D exhibited a weaker preference compared to subtype B. In addition, subtypes A, C and D had significantly less integration in heterochromatin (e.g. *LADs*) and *SINEs* compared to subtype B, which likely reflects their higher integration into genes. Pairwise analyses of the different integration site profiles showed that subtypes C and D, and B and D, shared the most similarity to each other, and subtype A differed the most from the other subtypes (**Figure 4D**). This relates in part to the sequence distance between IN coding regions of subtypes A, B, C, and D, with B and D being most genetically similar (**Figure 4E and 4F**). However, it is important to stress that any slight differences in integration profiles could also relate to discreet single amino acid differences between subtypes, especially in the CTD domain and/or the LEDGF binding domain.

**Figure 4:**
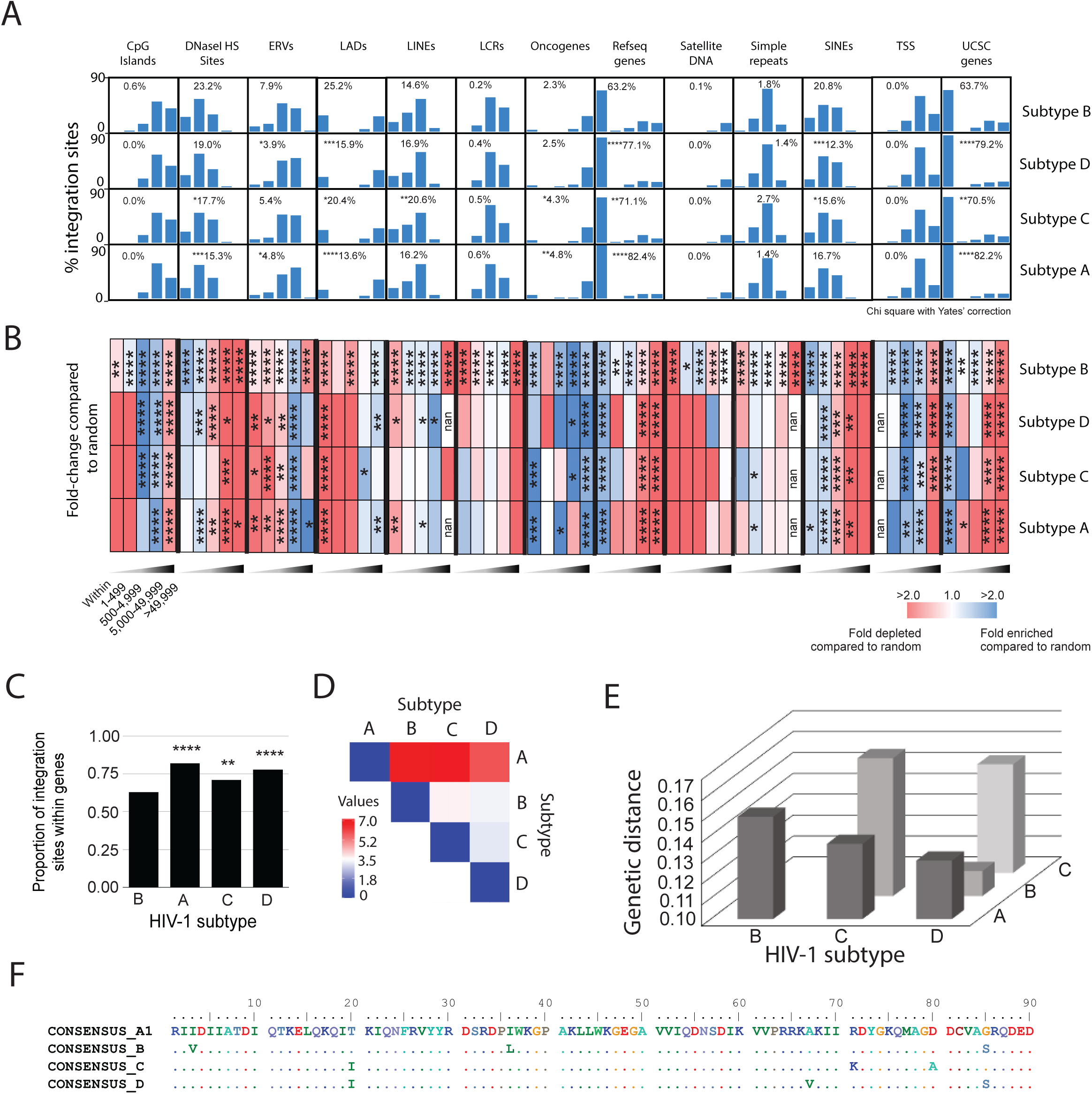
HIV-1 subtype A, B, C and D have different integration site preferences for genomic features. **A**, Comparison of the percentage of integration sites near common genomic features between HIV-1 subtypes A, B, C and D. **B**, Heatmaps depicting the fold enrichment or depletion of integration sites near common genomic features compared to the matched random control. Darker shades represent higher fold-changes in the ratio of integration sites to matched random control sites. Bins in A and B represent the distance of the integration sites from the genomic feature. **C**, Comparison of the percentage of integration sites within genes from HIV-1 subtypes A, B, C and D. **D**, Pairwise analysis was performed on the retroviral integration site profile preferences using Euclidean distance as the measurement method (Heatmapper, Babicki et al., 2016). Stronger relationships between retroviral integration site profiles are indicated by darker blue color in the pairwise distance matrix. Significant differences are denoted by asterisks *P < 0.05; **P < 0.01; ***P < 0.001; ****P < 0.0001 (Fisher’s Exact test). **E**, Estimates of Evolutionary Divergence over Sequence Pairs between Groups. The number of amino acid substitutions per site from averaging over all sequence pairs between groups are shown. Analyses were conducted using the Poisson correction model. This analysis involved 486 amino acid sequences. The coding data was translated assuming a Standard genetic code table. All ambiguous positions were removed for each sequence pair (pairwise deletion option). There were a total of 265 positions in the final dataset. Evolutionary analyses were conducted in MEGA X. **F**, Amino acid alignment of the C-terminal domain of HIV-1 integrase from subtypes A, B, C and D.

Analysis of the non-B DNA integration site profiles from the different subtype viruses showed enriched integration near non-B DNA (**Figure 5A, 5B and Table S7**). Notably, subtypes A and C had more integration sites near A-phased DNA compared to subtypes B and D. Subtypes A, C and D virus had significantly less sites near G4 DNA compared to subtype B virus. In addition, subtype C had significantly less sites near triplex and Z-DNA. Pairwise analysis of the different non-B DNA integration site profiles showed that the profiles of all subtypes differed substantially from each other, with subtypes B and D showing slightly more similarity to each other compared to the other subtypes (**Figure 5C**). Together, these data show that the integration site profiles from individuals infected with HIV-1 subtype B differs from those infected with non-subtype B virus, especially with respect to their preferences for non-B DNA. As discussed below, these slight but significant differences in selection for integration at different non-B DNA features may relate to discreet amino acid differences in the CTD domains of these IN that evolved in different subtypes. Although there are trends suggesting impact by discreet amino acid differences, for significance, we need to analyze more integration patterns by more subtype A, C, and D infections with known IN sequences.

**Figure 5:**
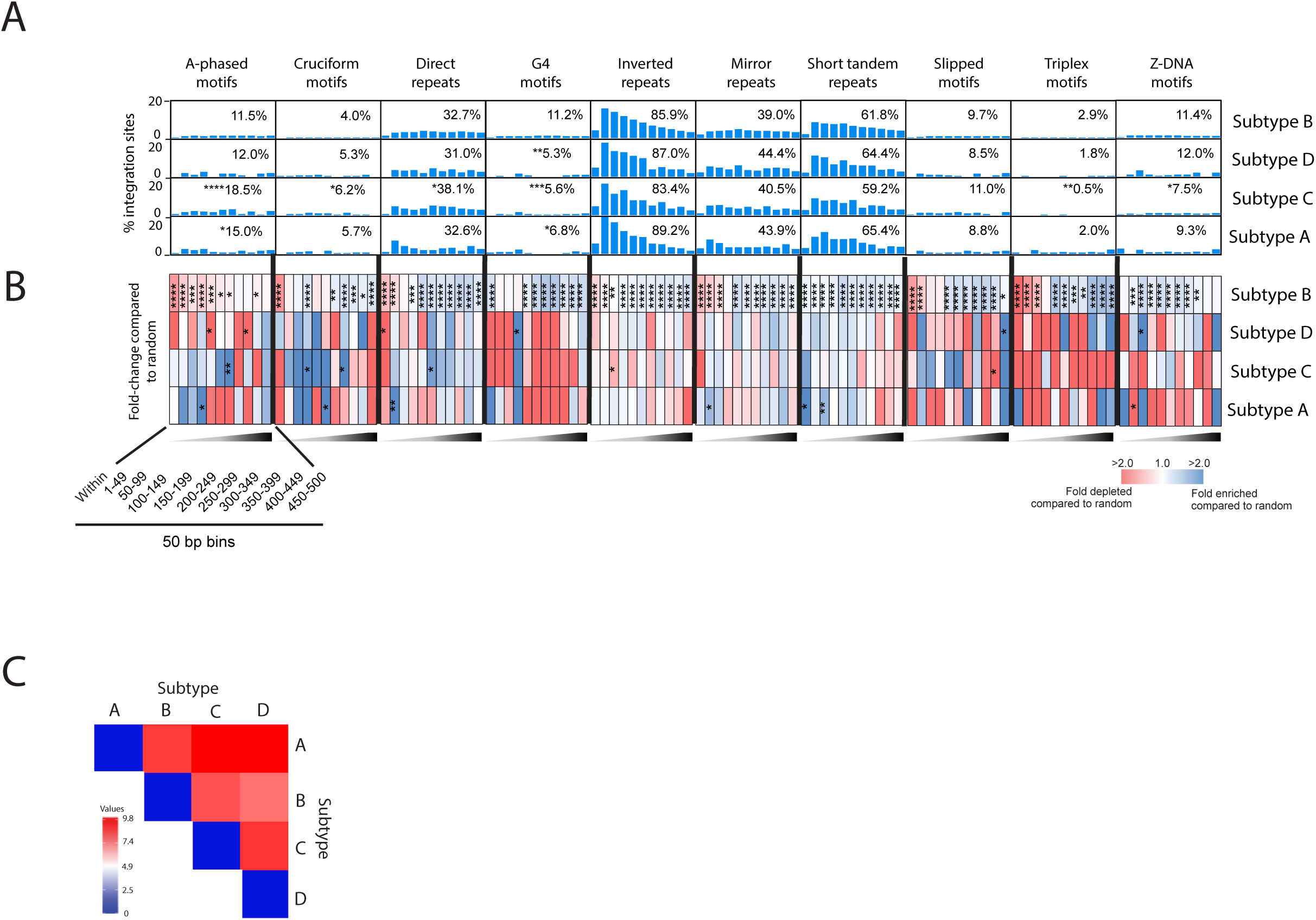
HIV-1 subtype A, B, C and D have different integration site preferences for non-B DNA. **A**, Comparison of the percentage of integration sites near non-B DNA features between HIV-1 subtypes A, B, C and D. **B**, Heatmaps depicting the fold enrichment or depletion of integration sites near non-B DNA compared to the matched random control. Darker shades represent higher fold-changes in the ratio of integration sites to matched random control sites. Bins in A and B represent the distance of the integration sites from the non-B DNA feature. Comparison of the percentage of integration sites within genes from HIV-1 subtypes A, B, C and D. **C**, Pairwise analysis was performed on the retroviral integration site profile preferences using Euclidean distance as the measurement method (Heatmapper, Babicki et al., 2016). Stronger relationships between retroviral integration site profiles are indicated by darker blue color in the pairwise distance matrix. Significant differences are denoted by asterisks *P < 0.05; **P < 0.01; ***P < 0.001; ****P < 0.0001 (Fisher’s Exact test).

### Integration hotspots are shared between HIV-1 subtypes

The concept of an HIV-1 integration “hotspot” was introduced to describe areas of the genome where integrations accumulate more than expected by chance in the absence of any selection process (*9, 61*). We catalogued and analyzed 1,000 bp windows containing two or more integration sites from each HIV-1 subtype dataset. This yielded a total of 10,892 hotspots between all subtypes (A: 41/429, 9.6%; B: 10,762/118,727, 9.1%; C: 74/484, 15.3%; D: 15/323, 4.6%; and MRC: 2/2982, 0.07%) (**Figure 6A**). 16% of the hotspots were shared between subtypes A and C, 14% between subtypes A and D and 10% between subtypes C and D. Despite the large number of hotspots identified in the subtype B dataset, less than 0.03% of the hotspots were shared between subtype B and the other subtypes. Comparison of hotspots revealed one hotspot in the *tbc1d5* gene that was shared by all subtypes (**Figure 6B and 6C**).

**Figure 6:**
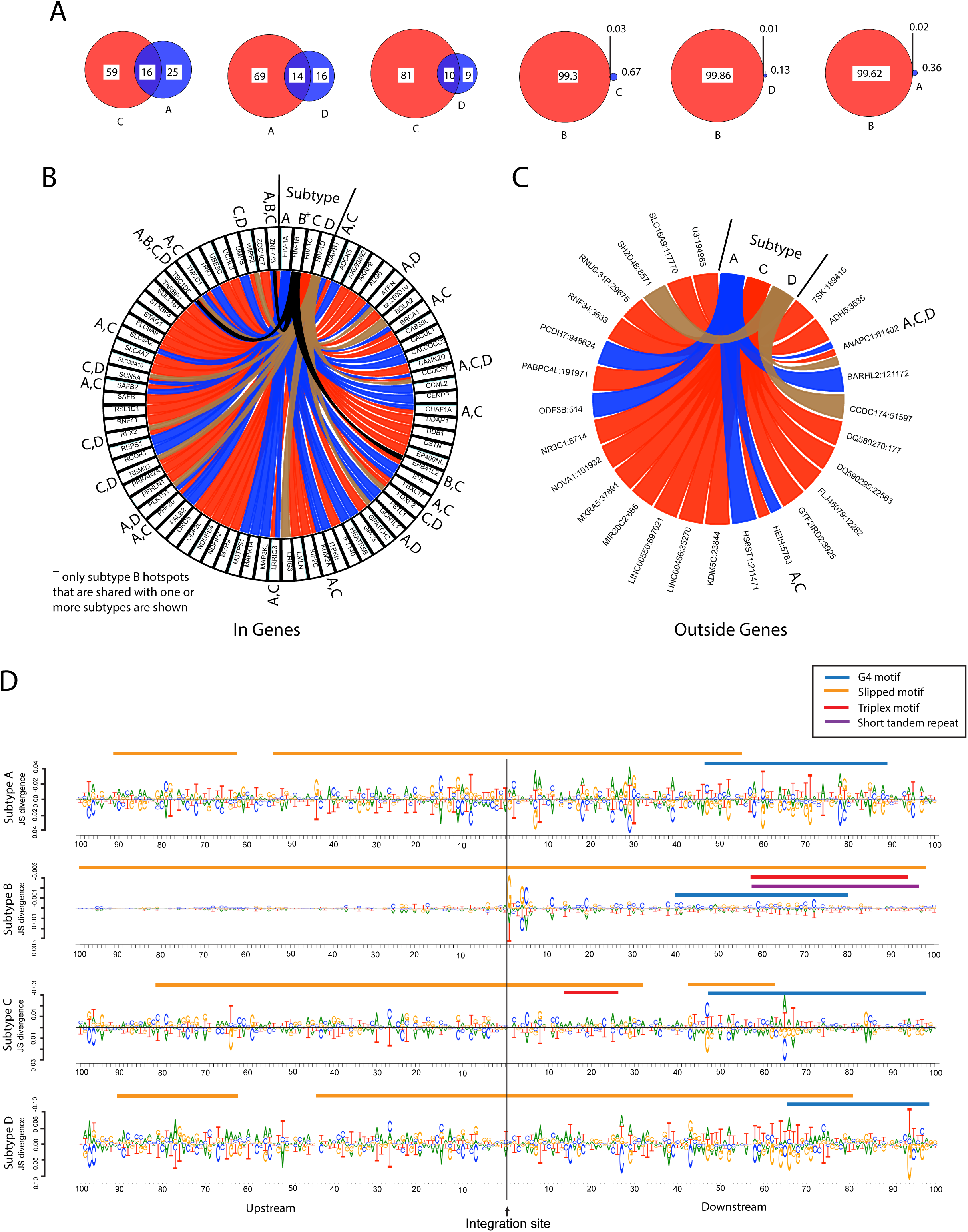
Integration hotspots identified from individuals infected with different HIV-1 subtypes. **A**, Venn diagram showing the percentage of unique (colored red or light blue) and shared (colored dark blue) integration site hotspots among the different HIV-1 subtypes. **B** and **C**, circos plots showing integration site hotspots inside (B) or outside (C) genes. The subtypes whose hotspots were shared is denoted in capital letters on the outside of the circos plots. For integration site hotspots outside genes (C), the closest gene and the distance from that gene are shown on the outside of the circos plots separated by a colon. Hotspots were defined as a 1 kb window in the genome hosting 2 or more integration sites. Note: (+) Due to the large number of subtype B integration site hotspots (10,762), only those hotspots shared between subtypes A, C and/or D are shown on the circos plots. No integration site hotspots outside of genes were shared between subtype B and any of the other subtypes. **D**, Consensus sequences were generated from a window of 100 nucleotides upstream and 100 nucleotides downstream of each integration site. Consensus sequences from integration sites located in hotspots were compared to consensus sequences from sites not located in hotspots using DiffLogo. Consensus sequences were analyzed for the presence of non-B DNA motifs (solid colored lines).

To identify potential local genomic sequences that may serve as ‘beacons’ for increased integration, we analyzed a nucleotide window of 100 bp upstream and 100 bp downstream of each integration site and generated a consensus sequence using DiffLogo (**Figure 6D**) (*62*). Comparison of the sequence motifs surrounding integration sites located in hotspots with the sequence motifs from sites not located in hotspots revealed the presence of a slipped DNA motif at the integration site in each of the HIV-1 subtype datasets. In addition, a G4 motif was located ∼40-100 bp downstream from the integration site in each of the different HIV-1 subtype datasets. Subtypes B and C also had a triplex motif and subtype B had a short tandem repeat located downstream of the integration site. Together, these data show that sites located in integration hotspot regions are located in slipped DNA motifs flanked downstream by other non-B DNA motifs.

## DISCUSSION

Here we showed that evolutionarily diverse retroviruses from all seven genera exhibit strong preferences for integration within or near non-B DNA features. Whereas all lentiviruses and most retroviruses integrate within or near genes and a preference for non-B DNA, we show that MMTV and ERV integration sites are highly enriched in heterochromatin but with still a preference for non-B DNA. Nonetheless, each retrovirus has distinct preferences for integration at specific types of non-B DNA. Highly distinct preferences for these non-B motifs among different HIV-1 subtypes is less apparent and yet, divergent IN evolution may still result in some differential selection/inhibition of HIV DNA integration in host cellular DNA. Finally, we showed that certain regions of the genome attract more integration sites than others and that the integration sites in these regions are located in slipped DNA motifs and flanked downstream by other non-B DNA motifs.

All retroviruses exhibited enriched integration near non-B DNA with specific non-B DNA features (e.g. slipped, G4, triplex and short-tandem repeat DNA) serving as integration site hotspots. How retroviral PICs recognize non-B DNA for integration is not fully understood. Non-B DNA motifs are enriched near endogenous retroviruses, are near sites of genomic variability, and are targets for preferential homologous recombination (>20-fold) in human cells (*63*–*65*). It is possible that integrase itself, components of the PIC and/or host proteins can bind non-B DNA directly and promote integration in these regions. We recently showed that PIC-binding host factors APOBEC3 (*39*), LEDGF/p75 and CPSF6 (submitted) influence the distribution of integration sites near non-B DNA features, potentially indicating that a host-directed mechanism is involved. Intriguingly, APOBEC3G binding and deamination hotspots also comprise slipped DNA motifs (*66, 67*). It is possible that viral integrase is also involved in recognizing non-B DNA given that HIV-1 integrase has been shown to bind directly to G4 DNA (*68*).

Analysis of integration site profiles after acute infection showed that HIV-1, FIV and SIV_mne_ had the strongest preference for integrating into transcriptionally active regions of the genome, whereas MMTV, FV and ERVs had the strongest preference for transcriptionally inactive regions. The targeting of transcriptionally active regions of the genome is believed to help maximize provirus expression for establishing infection and promoting its spread (*9*). Over time, especially with the help of antiretroviral drugs, the immune system likely clears these high virus-producing cells leaving behind infected cells that produce little to no virus that escape detection by the immune system. The integration profile of these latently infected cells showed that sites are located in more transcriptionally silent regions of the genome, similar to the profile shown here for ERVs and after acute infection with MMTV and FV. Consistent with the latter observation, latently infected cells have reduced integration in genes and increased integration in *SINEs* and heterochromatin, typically found in gene deserts or transcriptionally silent regions (*57, 60*). Transcriptionally silent regions of the genome are characterized by several features. For example, *LADs* represent a repressive chromatin environment tightly associated with the nuclear periphery (*56, 57*). *SINEs* (e.g. Alu repeats) and other transposed sequences are known to serve as direct silencers of gene expression due to their repressed chromatin marks (histone H3 methylated at Lys 9) (*69, 70*). Moreover, non-B DNA structures such as G4, cruciform, triplex and Z-DNA have been shown to silence expression of adjacent genes (*65, 71, 80, 72*–*79*). The ability to target silent regions of the genome for integration could be a long-term survival mechanism for both retroviruses and their hosts. By integrating into transcriptionally silent regions of the genome, these viruses can minimize their expression and avoid detection by the immune system. As an extreme example, endogenous retroviruses, some of which are ancient viruses that have been co-evolving with their hosts for hundreds of millions of years (e.g. ERVs and FV), are commonly integrated in transcriptionally silent regions of the genome away from genes (Figures 1 and 2) (*81*–*83*). The host also benefits with a lower risk of disease. This is seen with FV which persists in their primate hosts in the absence of disease, and MMTV which typically does not cause cancer unless it integrates near an oncogene (*84*). It is currently unknown if integration in transcriptionally silent regions of the genome is driven by retroviruses, the host or both. Interestingly, host APOBEC3G and APOBEC3F appear to promote a transcriptionally silent integration site profile of HIV-1, suggesting that the host may indeed contribute to this silent phenotype (*39*).

Genomic position effects have been shown to influence HIV-1 expression, which can ultimately impact disease progression (*57, 85*–*87*). As described above, HIV-1 infection of a host cell results in preferential integration in actively transcribing genes, due in part to reduced chromatin structure. Thus, it is not surprising to observe this finding following *in vitro* infections. However, even *in vitro*, there is evidence of preferential integration near non-B DNA motifs. Within a human host, enhanced HIV-1 integration within genes remains evident but now, there is greater enrichment of those exact integration sites being near transcription-silencing non-B DNA motifs, suggesting an enrichment in the pool of HIV-1 proviral DNA within transcriptionally silent sites in the genome. As described above, this may be the consequence of the immune system consistently clearing away cells propagating HIV-1 while the cells with silent integrants are slowly enriched. HIV-1 integration sites from these *in vitro* experiments were typically analyzed from cells infected for 2-3 days, whereas the *in vivo*-derived integration sites were obtained from chronically infected individuals whose cells contain integrated proviruses that have undergone more than 2 months of selection in the host. Consistent with previous findings (*57, 60*), HIV integrants were found enriched within features associated with reduced expression such as *LADs*, satellite DNA and *SINEs* when compared to *in vitro*-derived sites. Moreover, we identified a sequence-based integration site bias towards non-B DNA, particularly transcription silencing G4, triplex and Z-DNA.

Much of our knowledge of HIV-1 integration site selection has come from studies using subtype B virus. Our analyses showed that HIV-1 non-subtype B viruses have a much stronger preference for integrating into transcriptionally active regions of the genome compared to subtype B virus. This was characterized by increased integration in genes and decreased integration in genomic features associated with transcriptional silencing such as *LADs, SINEs*, G4, triplex and Z-DNA. Despite the enriched proviral integration into non-B DNA among all HIV-1 subtypes, there was some integration site preferences that differed between each of the HIV-1 subtypes. In the integrase coding region, amino acid differences between subtypes are as a high as 16% (subtype B vs. C) but typically less than 8% IN diversity among HIV-1 isolates of one subtype (*13, 88, 89*). This level of amino acid diversity is the highest among the enzymes encoded by the HIV-1 *pol* gene. The greatest diversity within HIV-1 IN is found in the CTD, which is involved in genomic DNA binding. Thus, it is reasonable to suspect that HIV-1 subtypes may target different non-B DNA motifs with differential selectivity. Despite the multiple substitutions that segregate HIV-1 variants of each subtype, there may be specific amino acid differences in one subtype versus another that impact preference for integration sites. For example, the isoleucine found at position 20 in subtype C and D versus the threonine in subtype A and B in the IN CTD may relate the greater similarity of integration site selection of subtypes C and D (Figure 4E and 4F). Despite the closer genetic relationship between HIV-1 subtype B and D across the genome, similar genetic distances separating subtypes A, B, C, and D are within the IN coding region. Polymorphisms in HIV-1 integrase have been reported that retarget integration away from gene dense regions, which also correlated with increased disease progression and virulence (*87*). This suggests that characteristics of integrase itself is a driver of integration site selection in retroviruses. We are currently exploring if subtype-specific polymorphisms in integrase may account for differences in the targeting of transcriptionally silent regions of the genome and of non-B DNA.

In conclusion, we identified non-B DNA as a feature surrounding integration sites that are targeted differentially by evolutionarily diverse retroviruses. We have also presented the first comparative look at the integration site profiles of several HIV-1 non-subtype B viruses and showed that they differed from HIV-1 subtype B profiles. Together, this data highlights important similarities and differences in retroviral integration site targeting that can be used in future studies to better understand the evolution of retroviral integration site targeting and how these viruses integrate into our genomes for long-term survival.

